# Hepatoprotective Effects of Vernonia amygdalina Leaf Extract: In Vivo and In Silico Evidence Against Ethanol-Induced Liver Injury

**DOI:** 10.1101/2025.10.08.678638

**Authors:** Kwadwo Fosu, Emmanuel Owusu Ansah, Francis Obeng, David Annor, Daniel Tei Adjirakor Mati, Jessica Mildred Ehiobu, Hijiratu Mustapha, Michella Naa Awula Atswei Laryea, Melissa Aikins, Ernest Ninson, Emmanuella Amakuor Apeku, Emmanuel Halm, Yakubu Adam, Prince Amoah Barnie, Foster Kyei

## Abstract

Alcoholic liver disease (ALD) remains a major global health burden with limited therapeutic options. In this study, we investigated the hepatoprotective effects of *Vernonia amygdalina* (VA) leaf extract against ethanol-induced liver injury using integrated in vivo and in silico approaches. Wistar rats were exposed to ethanol to induce liver damage, followed by treatment with VA extract (100-300 mg/kg) or silymarin (50 mg/kg). Liver function indices were evaluated through biochemical analysis. Network pharmacology analysis was performed to identify active compounds and their protein targets, and molecular docking assessed the binding affinities of key compounds. Ethanol exposure markedly increased serum SGOT, SGPT, ALKP, and total bilirubin while decreasing total protein and albumin. VA treatment significantly corrected these alterations in a dose-dependent manner. At 300 mg/kg, VA reduced SGPT from 110.31 to 88.65 U/mL, ALKP from 52.20 to 28.14 U/mL, and total bilirubin from 1.19 to 0.59 g/dL. Network pharmacology revealed flavonoids and sesquiterpene lactones as key active compounds targeting ALD-relevant proteins, including AKT1, CASP3, and HSP90AA1. Molecular docking demonstrated strong binding affinities (−9.5 kcal/mol) of cryptolepine, luteolin, and apigenin with AKT1 and HSP90AA1. VA provides both biochemical and mechanistic protection against ethanol-induced liver damage, highlighting its therapeutic potential in ALD management.

## INTRODUCTION

The liver plays a central role in maintaining systemic homeostasis by regulating nutrient metabolism, detoxifying xenobiotics, and supporting immune responses **[1]**. However, its constant exposure to both endogenous and exogenous compounds make it highly susceptible to toxic injury. Among liver disorders, alcoholic liver disease (ALD) is a major global health challenge, affecting over 3.3 million people in 2021 and accounting for nearly half of cirrhosis-related deaths **[2, 3]**. ALD progresses through stages ranging from steatosis to hepatitis, fibrosis, cirrhosis, and hepatocellular carcinoma **[4]**, driven by oxidative stress, mitochondrial dysfunction, lipid peroxidation, and pro-inflammatory signaling **[5]**.

Despite public health campaigns and clinical advances, ALD treatment remains limited, particularly in advanced disease. Current pharmacological agents, such as corticosteroids and antioxidants, offer modest benefits and carry risks including immune suppression, infection susceptibility, and poor long-term outcomes **[6]**. These challenges are further magnified in low-resource settings where access to costly treatments is restricted. Consequently, interest in medicinal plants as affordable, multi-target therapeutic options has grown. According to the World Health Organization, up to 80% of populations in developing countries rely on traditional medicinal plants for primary healthcare **[7]**.

*Vernonia amygdalina* (VA), commonly called bitter leaf, is a widely used African medicinal plant traditionally employed for malaria, gastrointestinal disorders, fevers, and liver ailments [**8**]. Its ethnomedical use is supported by observations of wild chimpanzees consuming the leaves for self-medication [**9**]. Phytochemical profiling has identified flavonoids (luteolin, apigenin), sesquiterpene lactones (vernodalin, vernolide), vernoniosides, and phenolic acids—compounds linked to antioxidant, anti-inflammatory, and hepatoprotective activities [**8**]. Preclinical studies have demonstrated VA’s protective effects against acetaminophen- and carbon tetrachloride-induced liver injury through modulation of oxidative stress and inflammation [**10, 11**].

However, despite this growing body of evidence, the molecular mechanisms by which VA mitigates ALD remain poorly defined. In this study, we combined in vivo ethanol-induced liver injury modeling with in silico network pharmacology, ADME profiling, and molecular docking to systematically elucidate VA’s bioactive compounds, target proteins, and potential pathways. This integrative approach enables mechanistic insights while supporting the principles of the 3Rs in animal research by reducing reliance on in vivo testing during early-stage drug discovery [**12, 13**]. We hypothesized that VA flavonoids and sesquiterpene lactones act on multiple liver-relevant proteins involved in oxidative stress, inflammation, and apoptosis, thereby conferring hepatoprotection in ALD.

**Figure 1.**
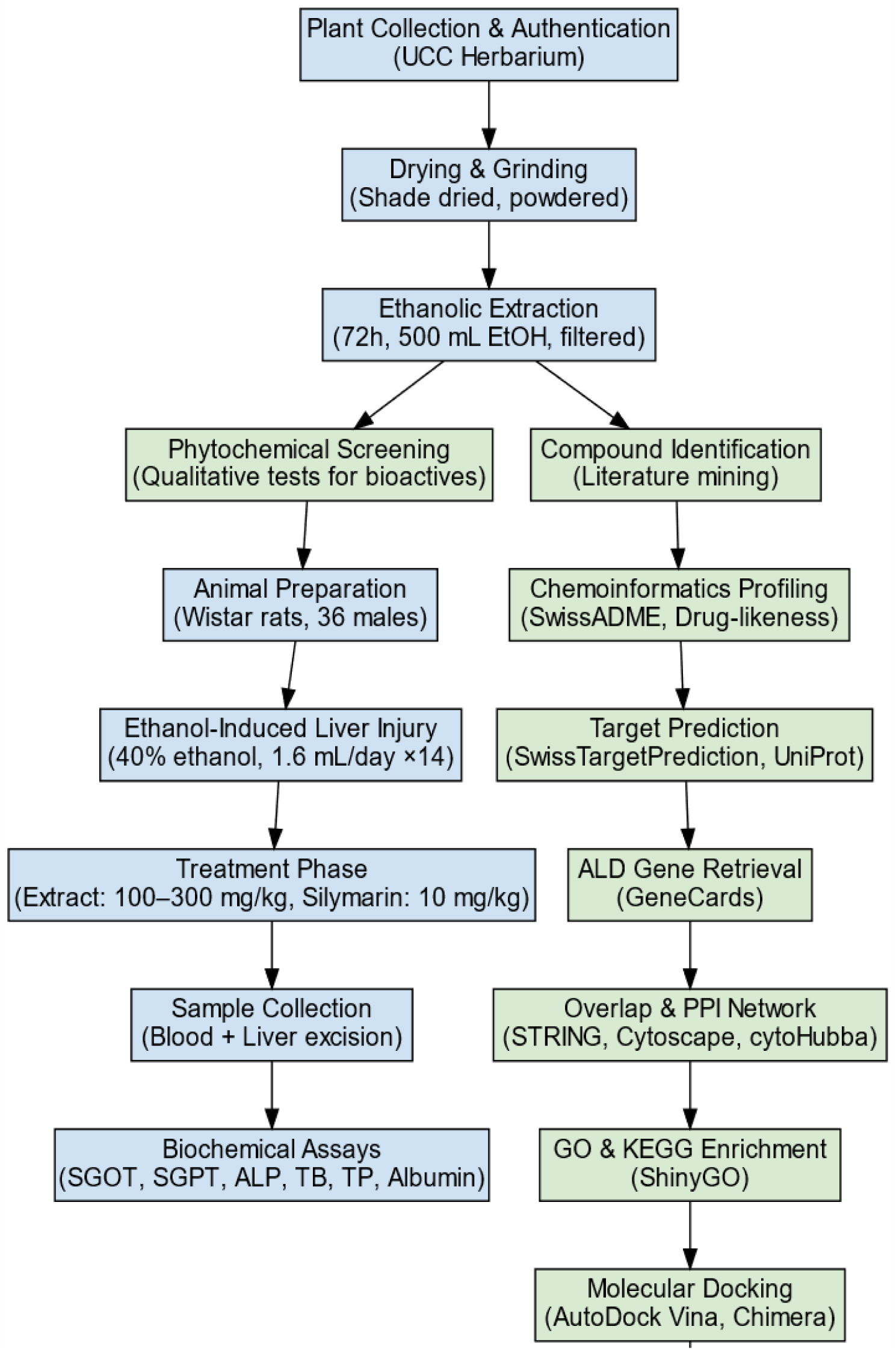
Workflow of in vivo and in silico studies on the hepatoprotective effects of *Vernonia amygdalina*.

## MATERIALS AND METHODS

### Plant Material Collection and Preparation

Fresh VA leaves were harvested from the University of Cape Coast campus, Ghana. The plant material was taxonomically authenticated at the Herbarium, School of Biological Sciences, University of Cape Coast, where a voucher specimen was deposited. Leaves were thoroughly rinsed with distilled water to remove surface debris, then air-dried in the shade at ambient room temperature (∼25°C) for three weeks. Dried leaves were ground into a fine powder using a mechanical blender and passed through a wire mesh sieve to ensure uniform particle size. The powdered material was stored in a sealed, labeled container at room temperature in a dry, dark environment until required for extraction and downstream analysis.

### Extraction, Filtration, and Concentration of Plant Material

Fifty grams of the powdered VA leaf sample were weighed into a 1 L flask and macerated in 500 mL of ethanol (added in two portions to ensure full submersion). The flask was sealed with cotton wool, wrapped in aluminum foil to minimize solvent loss, and placed on a shaker (Stuart Scientific, UK) at 250 rpm for 72 hours at room temperature. The mixture was then filtered sequentially through cotton wool and Whatman No. 1 filter paper. The filtrate was concentrated in a water bath at 37 °C to yield a semi-solid crude extract, which was allowed to cool, wrapped in foil, and stored in a desiccator containing activated silica gel until further use.

### Phytochemical Screening

To evaluate the phytochemical constituents of VA, a working solution was prepared by dissolving 0.5 g of the crude ethanolic extract in 50 mL of distilled water to obtain a 0.01 mg/mL solution. The solution was thoroughly shaken to ensure uniform dissolution and used directly for qualitative screening. Phytochemical screening was carried out by standard colorimetric assays described by Trease & Evans (2009) and Savithramma et al. (2011) [**14, 15**], with slight modifications. Tests for saponins, alkaloids, steroids, phenols, tannins, flavonoids, glycosides, cardiac glycosides, carotenoids, and terpenoids were conducted based on characteristic color changes, foaming, or precipitate formation.

### In Vivo Hepatoprotective Assessment

Thirty-six healthy male Wistar albino rats, each weighing approximately 200 g, were obtained from the Animal Holding Facility of the University of Cape Coast and acclimatized for one week under controlled laboratory conditions. Animals were housed in polypropylene cages (six rats per cage) under a 12-hour light/dark cycle, with a constant temperature range of 23–25 °C and relative humidity between 55% and 65%. During this period, rats had unrestricted access to standard pellet chow and distilled water. Alcoholic liver damage was induced in all groups except the naïve control by oral administration of 40% ethanol (1.6 mL per rat per day) for 14 consecutive days, based on previous research [**16**]. Following induction, rats were randomly allocated to six groups (n=6 each). The naïve and negative control groups received distilled water. The positive control group was treated with silymarin (10 mg/kg), a standard hepatoprotective agent in ethanol injury models **[16]**. The remaining groups received the VA extract at doses of 100, 200, or 300 mg/kg, administered once daily for 14 days. Figure 2 shows a schematic diagram of the experimental design for animal treatment

**Figure 2.**
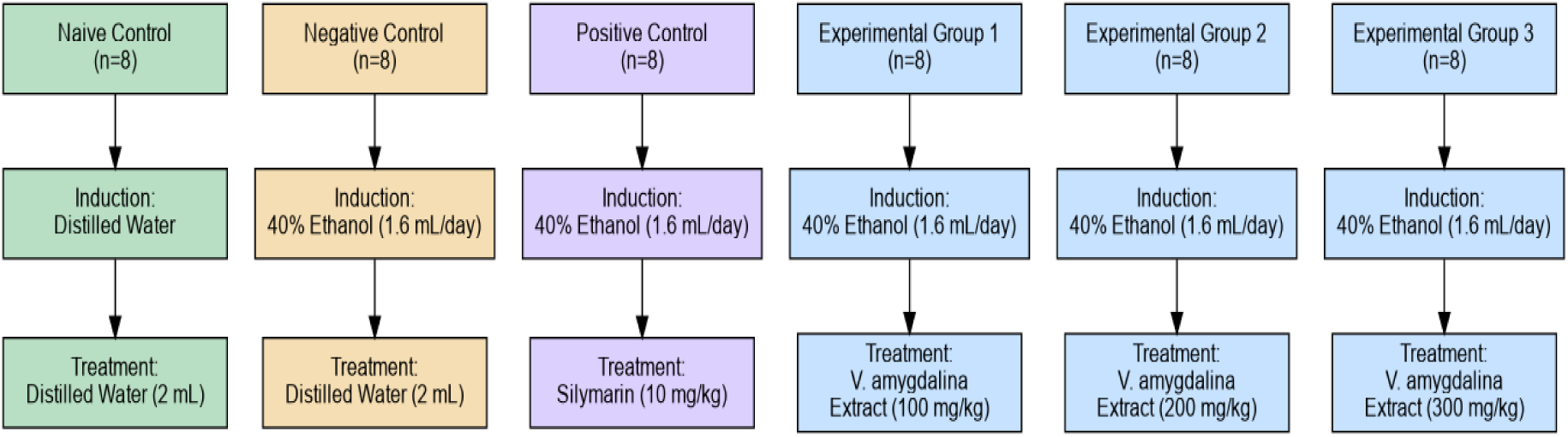
The experimental design for animal treatment

### Preparation of serum

Twenty-four hours after the final administration, one rat from each experimental group was euthanized. Euthanasia was carried out using either 2% isoflurane anesthesia or chloroform inhalation, following institutional ethical guidelines. Blood samples were obtained via cardiac puncture and immediately transferred into heparinized tubes. Samples were then centrifuged at 4,000 rpm for 15 minutes at 4 °C to separate the serum, which was subsequently stored at −20 °C until analysis.

### Enzyme Assays

Serum samples were sent to MKS Medical Laboratories (Spintex Road, Lashibi, Tema) for biochemical profiling. Liver function markers, including serum glutamate oxaloacetate transaminase (SGOT), serum glutamate pyruvate transaminase (SGPT), alkaline phosphatase (ALP), total bilirubin (TB), total protein (TP), and total albumin (TA), were quantified using the ABBOTT Afinion™ 2 Blood Analyzer (Abbott Laboratories, Chicago, IL, USA), following the manufacturer’s protocols.

### Statistical Analysis

All biochemical data were expressed as mean values with their corresponding standard errors of the mean (mean ± SEM). Statistical comparisons among treatment groups were performed using one-way analysis of variance (ANOVA) using Microsoft Excel 2019 MSO (Version 2505 Build 16.0.18827.20102), followed by Dunnett’s post hoc test to evaluate significant differences relative to the control group. A p-value of less than 0.05 or 0.01 was considered statistically significant. For the heatmap, data were normalized to the control group (set at 100% for each biomarker). Relative percentage values were calculated by dividing each group mean by the corresponding control mean and multiplying by 100.

### Literature-Based Identification of Bioactive Compounds

To identify candidate bioactive compounds with hepatoprotective potential, a structured literature search was conducted using databases such as PubMed, ScienceDirect, Google Scholar, Wiley Online Library, and SpringerLink. Boolean operators and wildcard terms were applied to refine search results. Key phrases included:

"Vernonia amygdalina AND phytochem", "bitter leaf AND hepatoprotective", "Vernonia amygdalina AND antioxidant", "bitter leaf AND a*nti-inflammatory"*, and *"Vernonia amygdalina AND alcoholic liver disease"*. The search targeted studies reporting the isolation, identification, or pharmacological profiling of secondary metabolites from *V. amygdalina*, particularly those linked to liver function, oxidative stress, or inflammation. Titles and abstracts were screened for relevance, and full texts were reviewed to confirm the identity of the compound and its biological significance. Compounds that overlapped across multiple sources and were supported by experimental evidence for hepatoprotective activity, especially in the context of alcohol-induced liver injury, were selected. The chemical structures of selected compounds were retrieved from PubChem in SDF or SMILES format for use in downstream computational analyses [**17**].

### Chemoinformatics and Drug-Likeness Profiling

The selected compounds from VA were subjected to chemoinformatic profiling to determine their potential as drug candidates. A comprehensive assessment of physicochemical properties, pharmacokinetic behavior, and oral bioavailability was performed using the SwissADME platform (http://www.swissadme.ch)[**18**]. SMILES structural data were input into the tool to generate relevant descriptors. Drug-likeness was evaluated based on key parameters, including molecular weight, lipophilicity (LogP), topological polar surface area (TPSA), solubility, and the number of hydrogen bond donors, acceptors, and rotatable bonds. Compounds demonstrating favorable ADME (Absorption, Distribution, Metabolism, Excretion) and toxicity profiles with hepatic relevance were selected for further analysis.

### Target Prediction of Bioactive Compounds

The potential protein targets of the selected VA compounds were predicted using the SwissTargetPrediction web tool (http://www.swisstargetprediction.ch) [**19**]. The canonical SMILES strings of each compound were submitted to estimate likely human protein targets based on chemical similarity and known ligand–target associations. Only targets with the highest probability scores were considered. To ensure consistency across downstream analyses, all target proteins were mapped and annotated using the UniProt database (https://www.uniprot.org) **[20]**.

### Identification of ALD-Associated Targets

To compile gene targets implicated in ALD, we queried the GeneCards Human Gene Database (https://www.genecards.org) using the search term “alcoholic liver disease” [**21**]. The resulting gene set was filtered to retain entries with high relevance scores based on functional association and disease linkage. Predicted protein targets of VA compounds were then compared with the ALD-related gene set. The intersection between the two datasets was visualized using the online Venn diagram tool provided by the VIB Bioinformatics Core https://bioinformatics.psb.ugent.be/webtools/Venn/. Common targets within the overlap were identified and considered as potential molecular bases for the hepatoprotective effects of the selected compounds.

### GO and KEGG Pathway Enrichment Analysis

To explore the biological relevance of the overlapping targets, Gene Ontology (GO) and Kyoto Encyclopedia of Genes and Genomes (KEGG) enrichment analyses were conducted using ShinyGO v0.77 http://bioinformatics.sdstate.edu/go/ **[22]**. Enrichment was performed with a significance cutoff of p < 0.05. GO terms were classified under biological processes, molecular functions, and cellular components, while KEGG analysis identified key signaling and metabolic pathways associated with the target genes. Results were visualized using bar and bubble plots to highlight significantly enriched terms. These analyses provided insight into the functional roles and pathway-level mechanisms through which VA compounds may exert hepatoprotective effects in the context of alcoholic liver disease.

### Protein–Protein Interaction (PPI) Network Construction

To investigate the functional connectivity of the predicted targets, a protein–protein interaction (PPI) network was constructed using the STRING database (https://string-db.org/) with the organism parameter set to Homo sapiens [**23**]. A high-confidence interaction score threshold (> 0.9) was applied to ensure specificity in the network. The resulting interaction data were exported and visualized using Cytoscape v3.9.1 for network analysis **[24]**. To identify key regulatory nodes, the cytoHubba plugin was used to rank targets based on topological features, with hub proteins defined by high-degree connectivity **[25]**. These hub genes were retained for downstream molecular docking and dynamics simulations to further explore their potential as therapeutic targets in alcoholic liver disease.

To investigate the functional connectivity of the predicted targets, a protein–protein interaction (PPI) network was constructed using the STRING database (https://string-db.org/), with the organism set to Homo sapiens and a high-confidence interaction score threshold (> 0.9) to ensure specificity [**26**]. The interaction data were imported into Cytoscape v3.9.1 (https://cytoscape.org/) for visualization, selection of key hubs, and subnetwork analysis [Shannon et al.]. Network topology parameters, including node degree and clustering coefficient, were computed using the Network Analyzer tool to characterize the overall structure of the PPI network. Key hub proteins were identified using the cytoHubba plugin based on degree centrality. [**25].** The top 10 hub genes were selected and used to construct a focused high-confidence subnetwork. This subnetwork was subjected to GO and KEGG pathway enrichment analyses to explore the biological significance of these central regulators. Additionally, the top hub genes were evaluated for therapeutic relevance in ALD through molecular docking.

### Molecular Docking Analysis

Molecular docking was carried out to predict the binding affinity and interaction profiles between selected VA compounds and key hub proteins implicated in alcoholic liver disease. The 3D structures of bioactive compounds were retrieved from the PubChem database in SDF format and converted to PDB using PyMOL [**27**]. Ligands were then prepared in AutoDockTools v1.5.7 by removing water molecules, adding polar hydrogens, assigning Gasteiger charges, and saving them in PDBQT format after energy minimization using steepest descent optimization in UCSF Chimera v1.17 [**28, 29**]. Protein structures for docking were obtained from the RCSB Protein Data Bank [**30**], prioritizing high-resolution crystallographic models of targets identified from the PPI network. Co-crystallized ligands, ions, and water molecules were removed. Hydrogens and partial charges were added, and missing atoms were corrected using AutoDockTools. The final structures were saved in PDBQT format for docking. Docking simulations were performed using AutoDock Vina v1.1.2 [**31**], with each receptor prepared in a flexible docking grid box that encompassed the predicted or known active site region. Binding affinity scores (in kcal/mol) were recorded, and the best-ranked poses were analyzed further. Protein–ligand interactions were visualized using ChimeraX and BIOVIA Discovery Studio Visualizer to assess bond interactions [**32, 33**]. All methods employed in this study are summarized in **Figure. 1**.

### Ethical Considerations

All animal experiments were approved by the Institutional Animal Care and Use Committee (IACUC) of the University of Cape Coast.

## RESULTS

### Phytochemical Analysis

In this study, we aimed to evaluate the hepatoprotective potential of VA leaf extract using both in vivo and in silico approaches. In the primary analysis, qualitative phytochemical screening was conducted to identify bioactive constituents that may contribute to the therapeutic effect of the extract. The ethanolic extract revealed the presence of saponins, steroids, flavonoids, cardiac glycosides, glycosides, and terpenoids, while alkaloids, phenols, tannins, and carotenoids were absent **(Table 1)**. The presence of these secondary metabolites provides a biochemical basis for the subsequent pharmacological evaluation.

**Table 1.** Phytochemical constituents of ethanolic extract of Vernonia amygdalina leaves. The presence and quantity of each phytochemical were assessed using standard qualitative methods. (+++): strongly present; (---): absent.

| Phytochemical | Test Result | Inference |
| --- | --- | --- |
| Saponins | +++ | Present |
| Alkaloids | --- | Absent |
| Steroids | +++ | Present |
| Phenols and Tannins | --- | Absent |
| Flavonoids | +++ | Present |
| Cardiac Glycosides | +++ | Present |
| Glycosides | +++ | Present |
| Carotenoids | --- | Absent |
| Terpenoids | +++ | Present |

Ethanol exposure caused pronounced liver damage, evidenced by elevated serum SGOT, SGPT, ALKP, and TB levels, alongside decreased TP and TA levels relative to controls (**Figure 3**). Specifically, SGOT, SGPT, ALKP, and TB increased to 166.72 U/mL, 103.37 U/mL, 64.59 U/mL, and 2.34 g/dL, respectively, while TP and TA dropped to 6.86 g/dL and 3.19 g/dL. Treatment with VA extract (100–300 mg/kg) resulted in a clear, dose-dependent hepatoprotective response, significantly lowering transaminase and bilirubin levels while restoring TP and TA toward baseline values. The normalized heatmap (**Figure 4**) further illustrates this trend, highlighting the extract’s ability to reverse ethanol-induced alterations in liver biomarkers. Notably, at 300 mg/kg, VA restored hepatic parameters to near-control levels, comparable to the effect observed with silymarin, supporting its potential as an effective natural hepatoprotective agent. Raw means and statistical analyses are provided in Supplementary table 1.

**Figure 3.**
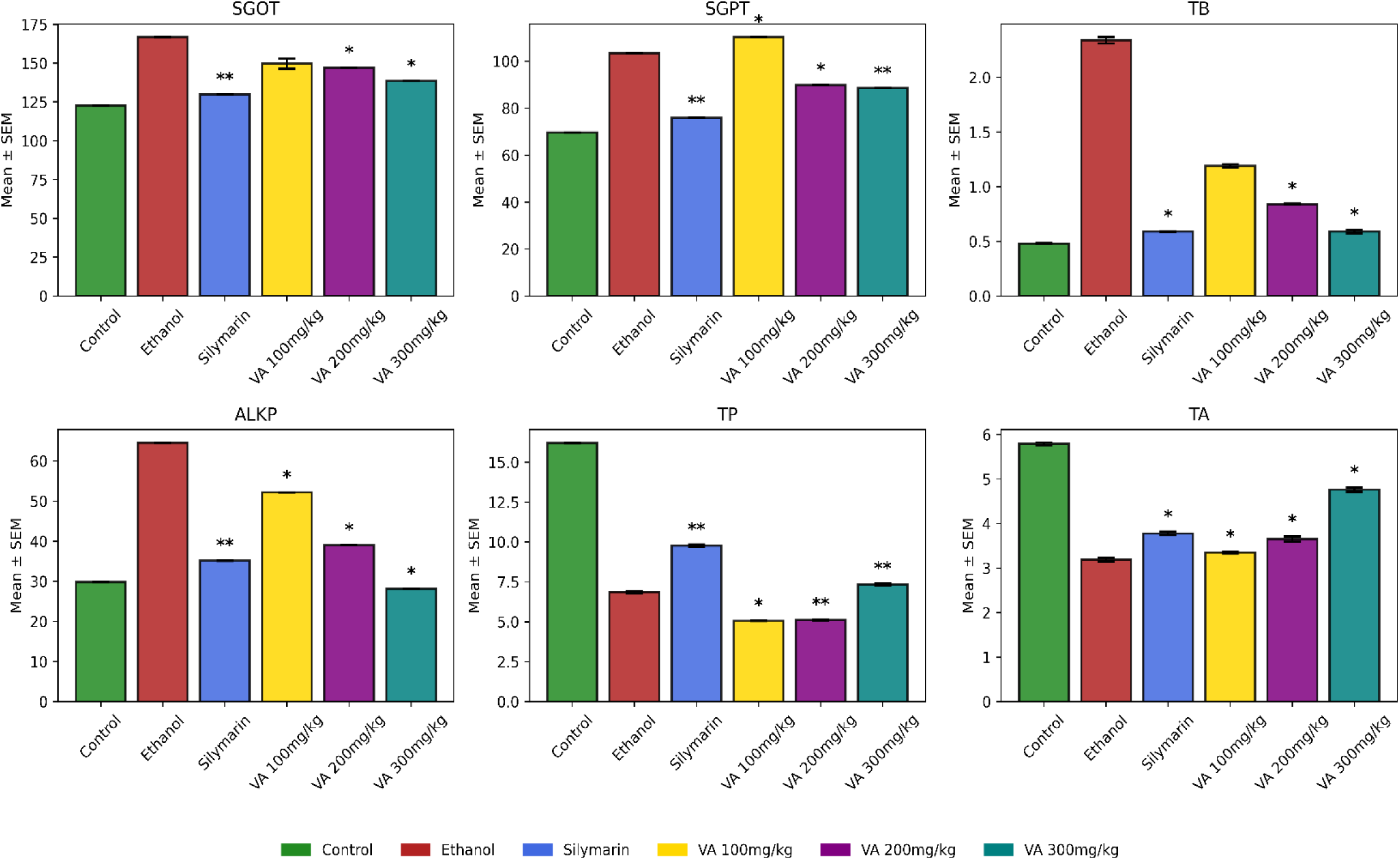
Effect of *Vernonia amygdalina* extract on liver biomarkers in ethanol-induced hepatotoxicity. Data are mean ± SEM (n = 6) for SGOT, SGPT, TB, ALKP, TP, and TA across groups: control, ethanol-only, silymarin (10 mg/kg), and VA (100–300 mg/kg). One-way ANOVA with Dunnett’s post hoc test; asterisks indicate significant differences vs. ethanol-only (*p* < 0.05; **p** < 0.01).

**Figure 4.**
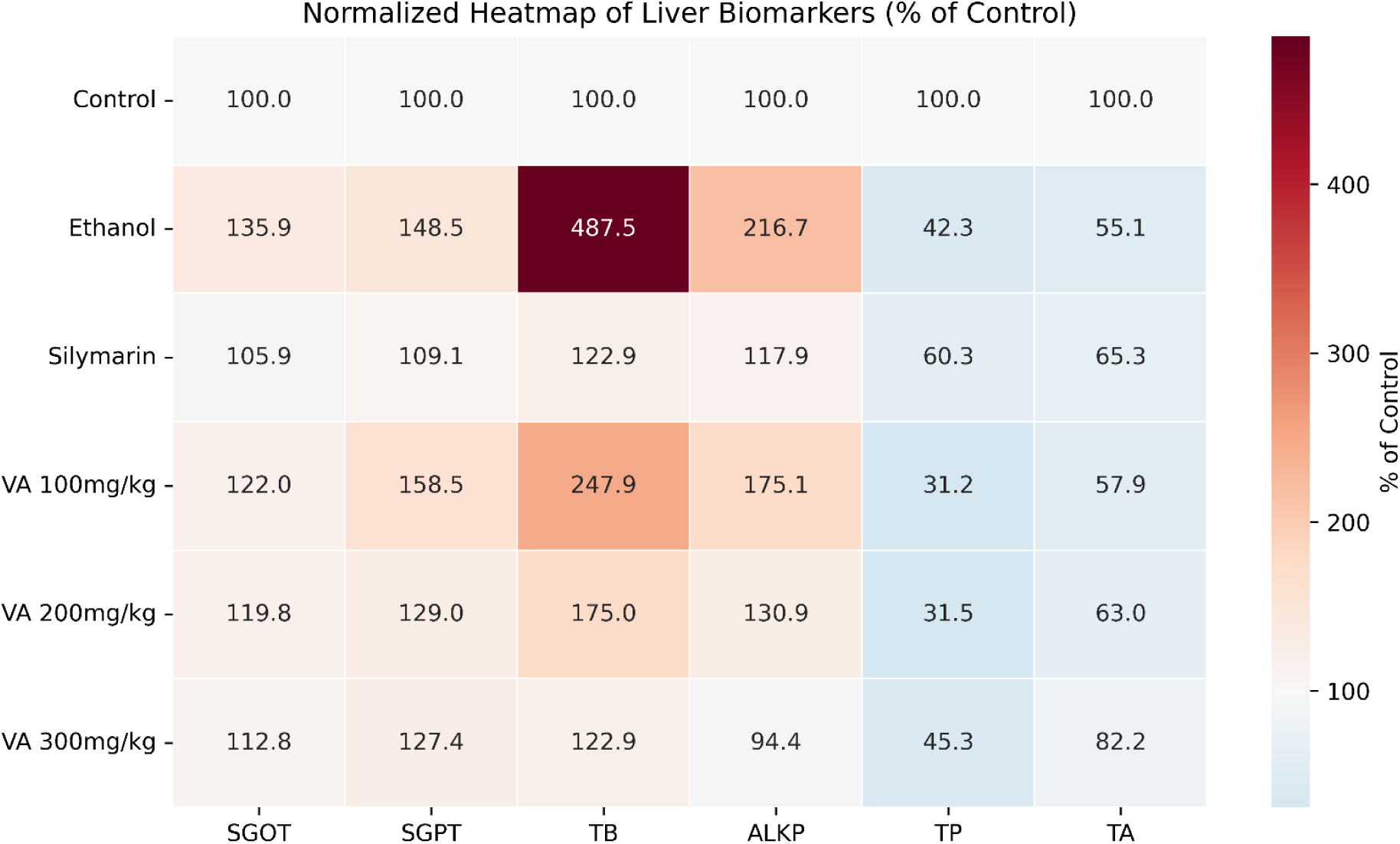
Heatmap representing the relative percentage change of liver function biomarkers in ethanol-induced hepatotoxic rats treated with VA extract. Data are normalized to the control group (set at 100% for each biomarker). Colors indicate relative increases (red) or decreases (blue) compared to the control.

### Literature-Based Identification of Bioactive Compounds

A structured literature search identified fifteen compounds reported in VA leaves with potential hepatoprotective, antioxidant, and anti-inflammatory activities. These include flavonoids (luteolin, apigenin, luteolin 7-*O*-glucoside), alkaloids (cepharanthine, cryptolepine), phytosterols (β-sitosterol, stigmasterol), sesquiterpene lactones (vernodalol, vernolide, hydroxyvernolide, dihydrovernodalin), and other secondary metabolites such as amygdalin, vernadolin, ingenol-3-angelate, and vernomenin. These phytochemicals are known to modulate oxidative stress and inflammation, supporting the observed hepatoprotective effects in ethanol-induced liver injury in rats. A summary of these compounds and their 3D structures is provided in Supplementary **Table S2**.

### Drug-likeness and Pharmacokinetic Screening

To identify active constituents of VA, the 15 phytochemicals were evaluated for drug-likeness using Lipinski’s Rule of Five and related oral bioavailability criteria **[34]**. Compounds with more than one violation, molecular weight ≥500 Da, logP outside the −0.7 to 5.0 range, hydrogen bond donors greater than five, acceptors greater than ten, or topological polar surface area (TPSA) above 140 Å² were excluded. Seven compounds, including luteolin, apigenin, cryptolepine, vernolide, dihydrovernodalin, hydroxyvernolide, and vernomenin, met all selection criteria (**Table 2**). Further filtering based on ADME properties showed that all seven compounds were predicted to have high gastrointestinal absorption and low blood–brain barrier permeability, suggesting activity limited to peripheral targets (**Table 3**)

**Table 2.**
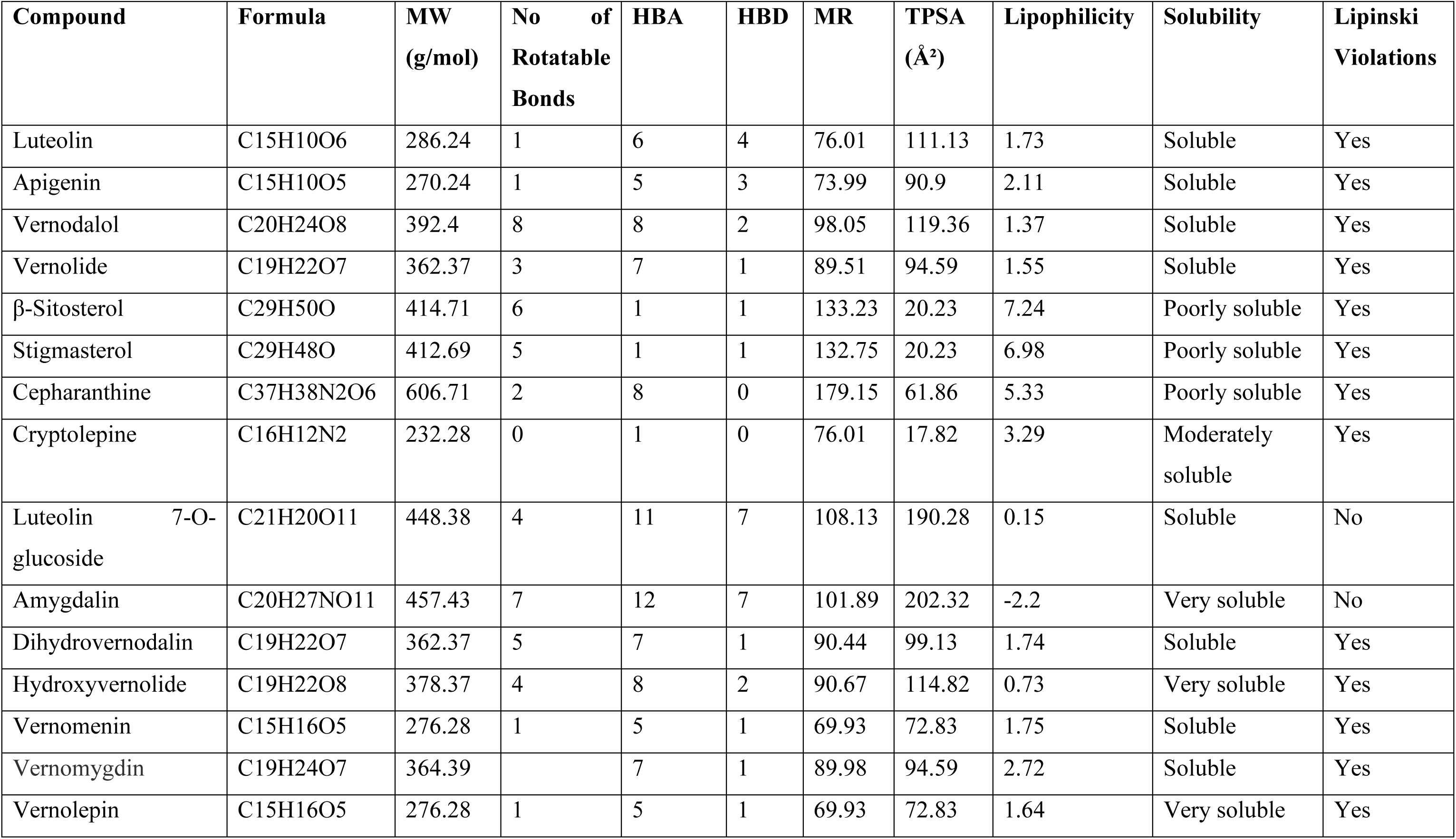
Physicochemical properties and drug-likeness of selected compounds from VA. MW: Molecular weight; HBA: Hydrogen bond acceptors; HBD: Hydrogen bond donors; MR: Molar refractivity; TPSA: Topological polar surface area. Except for cryptolepine, none of the selected compounds were predicted to be substrates of P-glycoprotein (P-gp), which minimizes the risk of efflux-mediated absorption loss. CYP450 profiling showed that luteolin, apigenin and cryptolepine may inhibit CYP1A2, CYP2D6 and CYP3A4 isoforms. In contrast, vernolide, hydroxyvernolide, dihydrovernodalin and vernomenin showed no inhibitory effects on the five major CYP isoforms, suggesting a lower risk of metabolic interactions. The final seven compounds were retained for downstream target prediction and molecular docking analysis.

**Table 3.** Pharmacokinetics and ADME prediction of compounds from Vernonia amygdalina. HIA: Human intestinal absorption; BBB: Blood–brain barrier permeability; P-gp: P-glycoprotein; CYP: Cytochrome P450.

| **Compound** | **GI Absorption** | **BBB Permeant** | **P-gp Substrate** | **CYP1A2 Inhibitor** | **CYP2C19 Inhibitor** | **CYP2C9 Inhibitor** | **CYP2D6 Inhibitor** | **CYP3A4 Inhibitor** | **Log Kp (cm/s)** |
| --- | --- | --- | --- | --- | --- | --- | --- | --- | --- |
| Luteolin | High | No | No | Yes | No | No | Yes | Yes | -6.25 |
| Apigenin | High | No | No | Yes | No | No | Yes | Yes | -5.8 |
| Vernodalol | High | No | No | No | No | No | No | No | -7.86 |
| Vernolide | High | No | No | No | No | No | No | No | -7.85 |
| β-Sitosterol | Low | No | No | No | No | No | No | No | -2.2 |
| Stigmasterol | Low | No | No | No | No | Yes | No | No | -2.74 |
| Cepharanthine | High | No | No | No | No | No | No | No | -5.36 |
| Cryptolepine | High | Yes | Yes | Yes | Yes | No | Yes | Yes | -5.35 |
| Luteolin 7-O-glucoside | Low | No | Yes | No | No | No | No | No | -8.00 |
| Amygdalin | Low | No | No | No | No | No | No | No | -11.01 |
| Dihydrovernodalin | High | No | No | No | No | No | No | No | -7.28 |
| Hydroxyvernolide | High | No | No | No | No | No | No | No | -8.84 |
| Vernomenin | High | No | No | No | No | No | No | No | -7.03 |
| Vernomygdin | High | No | No | No | No | No | No | No | -7.81 |
| vernolepin | High | No | No | No | No | No | No | No | -7.77 |

### Target identification and analysis

To identify potential molecular targets involved in the hepatoprotective effects of VA, an in-silico target prediction was performed for the seven shortlisted compounds using the SwissTargetPrediction platform (**Daina et al., 2019**). This process yielded 392 unique human protein targets. In parallel, 9,584 ALD-associated genes were retrieved from the GeneCards Human Gene Database (**Stelzer et al., 2016**), which integrates gene–disease associations from text mining, functional genomics, and pathway data. By intersecting the predicted targets of VA compounds with the ALD gene set, 351 common targets were identified (**Figure. 5A**). These overlapping genes may play a key role in mediating the hepatoprotective effects of VA phytochemicals and were employed in protein–protein interaction analysis.

**Figure 5.**
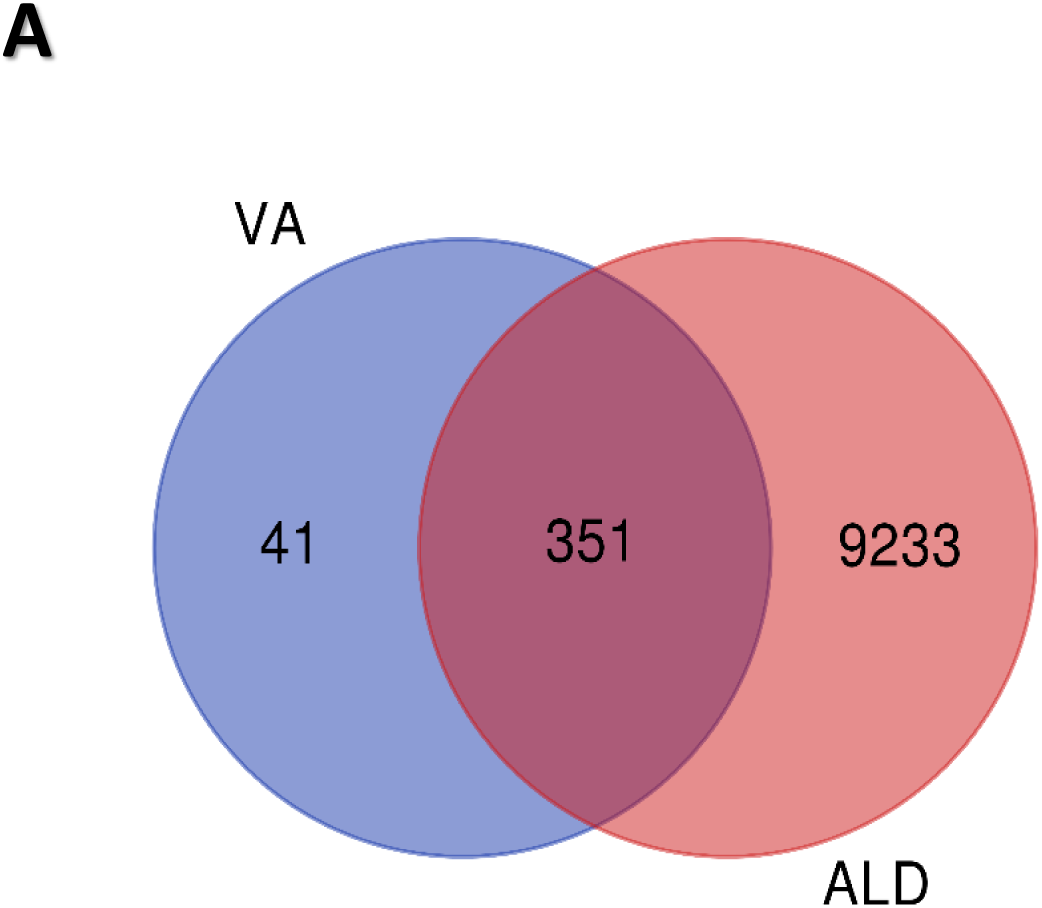
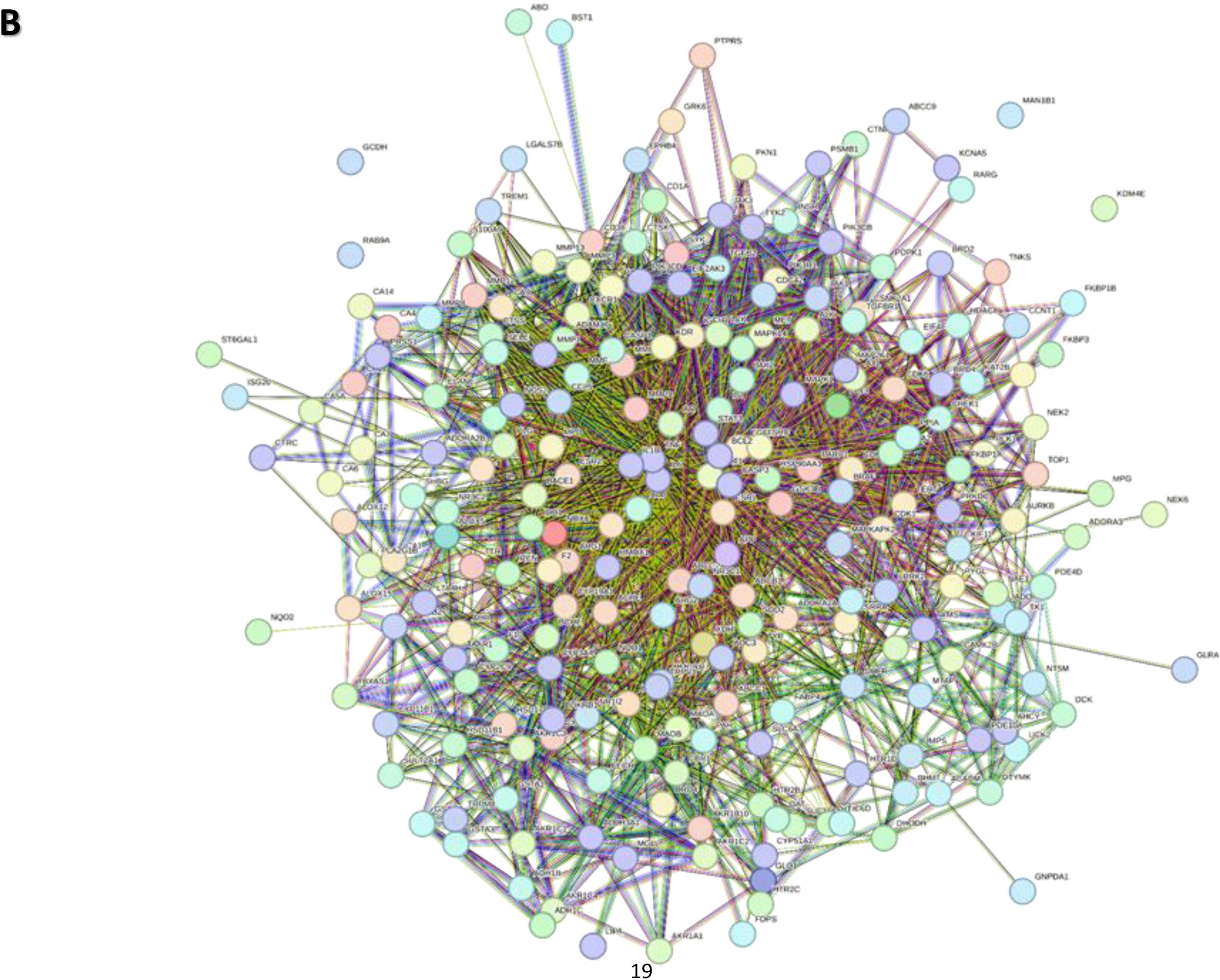
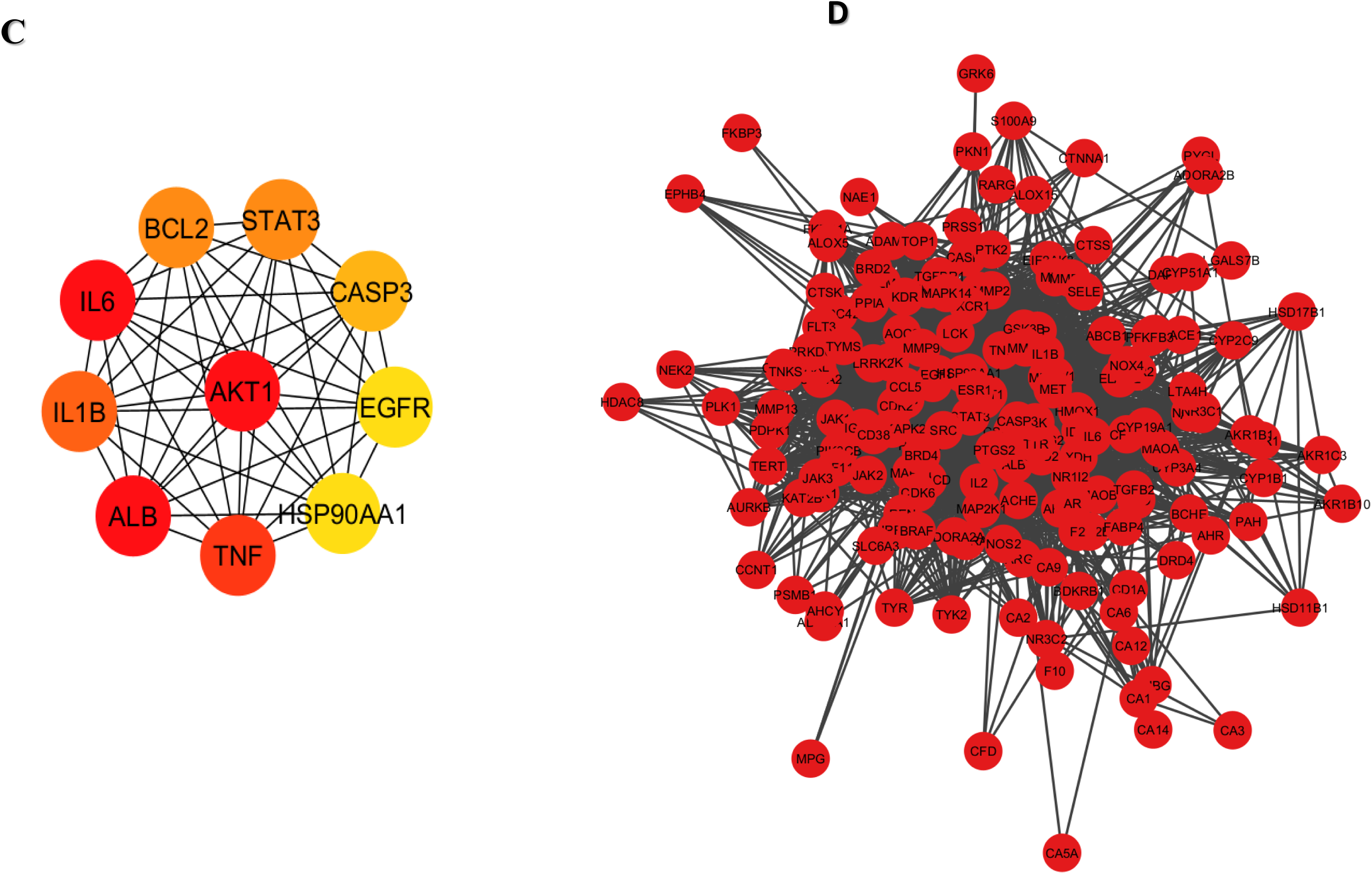
Protein–protein interaction (PPI) network and hub gene analysis: (A) Venn diagram showing the overlapping targets between *Vernonia amygdalina*-derived compounds and ALD-associated genes; (B) PPI network of the 351 overlapping targets constructed using the STRING database; (C) top 10 hub genes identified via the cytoHubba plugin in Cytoscape; and (D) subnetwork of key targets derived from the top hub genes.

**Table 5.** Top 10 Proteins in the STRING PPI Network Ranked by Degree Method.

| Rank | Gene Symbol | Protein Name | UniProt ID | Degree Score |
| --- | --- | --- | --- | --- |
| 1 | ALB | Serum albumin | P02768 | 112 |
| 1 | IL6 | Interleukin-6 | P05231 | 112 |
| 1 | AKT1 | RAC-alpha serine/threonine-protein kinase | P31749 | 112 |
| 4 | TNF | Tumor necrosis factor | P01375 | 111 |
| 5 | IL1B | Interleukin-1 beta | P01584 | 97 |
| 6 | STAT3 | Signal transducer and activator of transcription 3 | P40763 | 95 |
| 6 | BCL2 | B-cell lymphoma-2 | P10415 | 95 |
| 8 | CASP3 | Caspase-3 | P42574 | 92 |
| 9 | EGFR | Epidermal growth factor receptor | P00533 | 88 |
| 9 | HSP90AA1 | Heat shock protein HSP 90-alpha | P07900 | 88 |

### Network analysis of protein-protein interactions and identification of key targets

To understand how the differentially associated proteins interact, we used the STRING database (confidence score > 0.9; organism: *Homo sapiens*) to construct a protein-protein interaction (PPI) network. The network consisted of 233 nodes and 2,746 edges (**Figure. 5B**), where each node represents a protein and each edge denotes a predicted or experimentally validated interaction. The edges in the STRING network are supported by various types of evidence, including neighborhood, gene fusion, co-occurrence, experimental data, curated databases, text mining, and co-expression. In the evidence view, these edge types are represented by colored lines: green (neighborhood), red (fusion), blue (co-occurrence), purple (experimental), yellow (text mining), light blue (database), and black (co-expression). The dense presence of purple and green edges within the network indicates a strong basis in both experimental findings and neighborhood associations, suggesting reliable functional linkages between the proteins. The network was further visualized in Cytoscape (v3.9.1) using confidence mode, where the width of the edges corresponds to the confidence score of the interaction, with thicker lines denoting higher confidence. The overall network exhibited an average node degree of 23.6 and a local clustering coefficient of 0.539, indicating a densely interconnected topology. Using the cytoHubba plugin in Cytoscape, hub proteins were ranked by degree centrality to identify those with the highest number of interactions. The top 10 hub proteins were ALB, IL6, AKT1, TNF, IL1B, STAT3, BCL2, CASP3, EGFR, and HSP90AA1 (**Figure. 5C**), visualized with color intensity proportional to their degree values, with red indicating the highest centrality. These hubs are likely to play central roles in the underlying biological mechanisms of ALD. To refine the analysis, a high-confidence subnetwork consisting of the 10 hub proteins and their direct interactors was extracted. This subnetwork comprised 173 nodes and 2,406 edges (**Fig. 5D**), forming a single connected component. Topological analysis using Cytoscape’s Network Analyzer showed a network diameter of 6, characteristic path length of 1.990, clustering coefficient of 0.298, and network density of 0.081, with an average of 27.8 neighbors per node.

### GO and KEGG analysis

To understand the potential protective mechanisms of VA against ALD, we conducted GO and KEGG pathway enrichment analyses on 173 subnetwork targets derived from the top 10 hub genes identified in the PPI network. GO enrichment yielded a total of 1,835 significantly enriched terms (FDR < 0.05), comprising 1,000 biological processes (BP), 572 molecular functions (MF), and 263 cellular components (CC). The top BP were primarily associated with responses to nitrogen compounds, oxidative stress, small molecules, and the regulation of protein phosphorylation (**Figure. 6A**). In the MF category, enriched terms included ATP binding, protein kinase activity, notably serine/threonine and tyrosine kinases, and nucleotide binding (**Figure. 6B**). CC enrichment revealed strong associations with membrane rafts, extracellular vesicles, and synaptic membranes (**Figure. 6C**). KEGG pathway analysis identified 222 significantly enriched pathways (FDR < 0.05), highlighting PI3K–Akt, VEGF, HIF-1, FoxO, TNF, and IL-17 signaling, as well as lipid and atherosclerosis pathways (**Figure. 7**). These pathways are known regulators of inflammation, oxidative stress, angiogenesis, and metabolic homeostasis, which are hallmarks of liver injury and repair.

**Figure 6.**
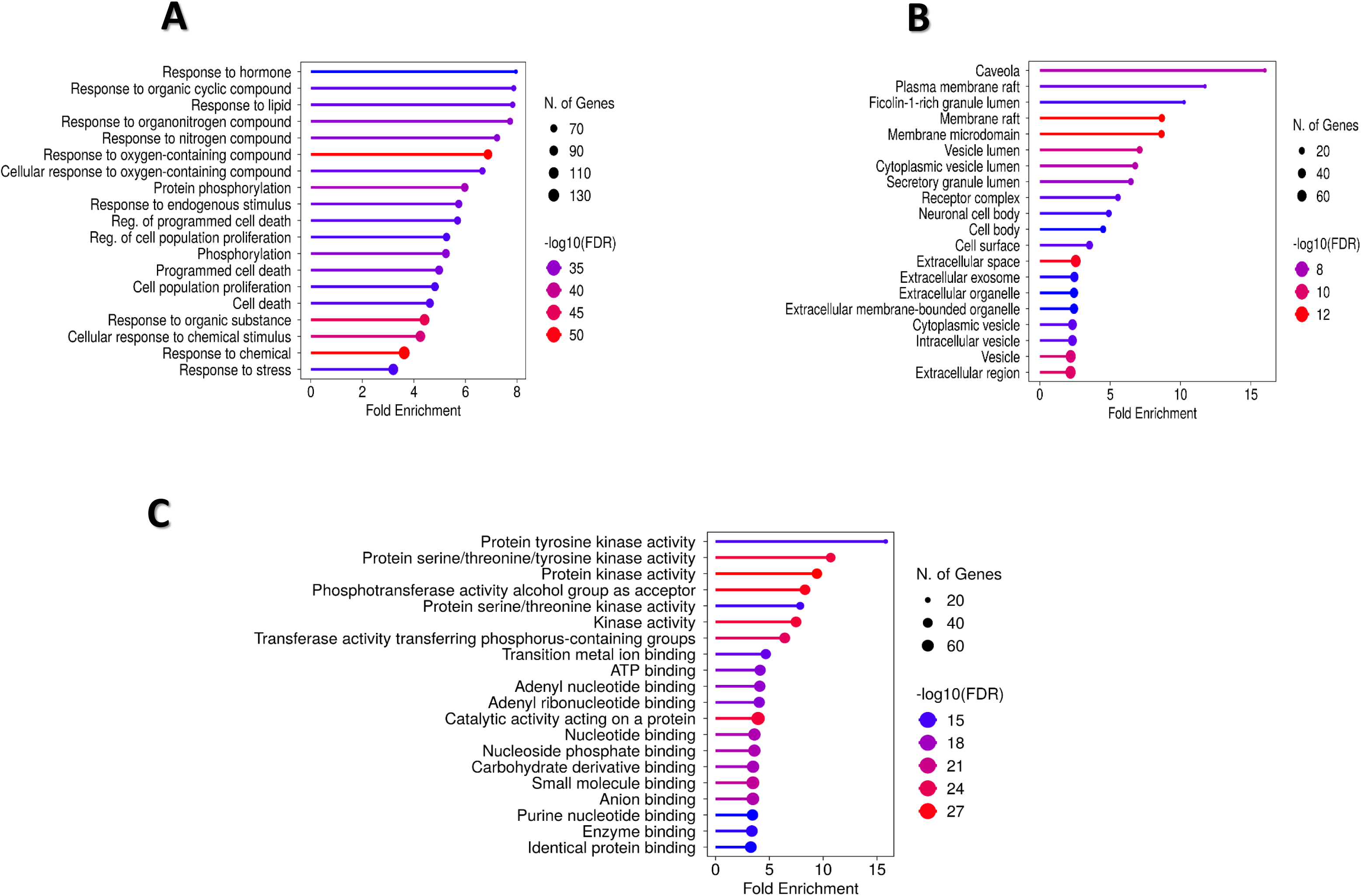
GO enrichment analysis of predicted targets of bioactive compounds from Vernonia amygdalina (VA) in ALD. Significant GO terms (p < 0.05) were identified for (A) biological processes, (B) cellular components, and (C) molecular functions related to the potential targets of VA in ALD

**Figure 7.**
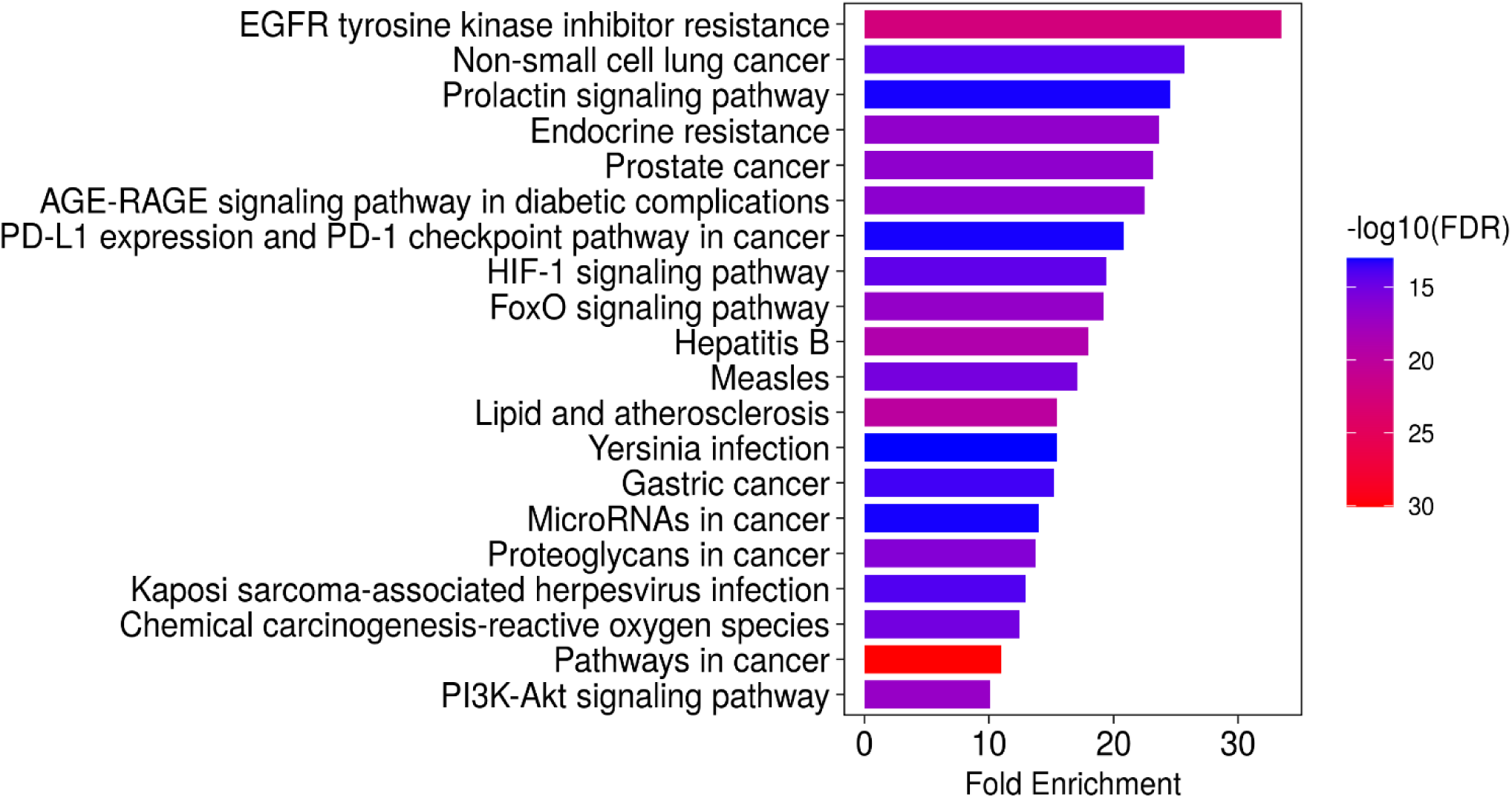
Top 20 KEGG pathways significantly enriched for the target genes associated with alcoholic liver disease

### Molecular Docking Analysis of Bioactive Compounds with Target Proteins

To validate the interactions between VA phytochemicals and key protein targets associated with ALD, molecular docking was carried out and the resulting binding affinities were evaluated. Compounds with binding energies of −9.5 kcal/mol or lower were considered to have strong binding potential. The docking results revealed binding energies ranging from −5.6 to −10.2 kcal/mol, with AKT1 and HSP90AA1 emerging as the most promising targets (**Table 6**). For the AKT1 protein (PDB: 3O96), cryptolepine demonstrated the highest affinity at −10.15 kcal/mol, followed by luteolin at −9.84 kcal/mol and apigenin at −9.52 kcal/mol (**Table 6**). Cryptolepine formed an electrostatic π-anion interaction with ASP292 (3.88 Å), π–π stacking with TRP80 (3.73 Å), and multiple hydrophobic contacts involving TRP80, LEU210, LYS268, VAL270, and TYR272 (**Figure 8C**). Luteolin **(Figure 8A)** and apigenin **(Figure 8B)** displayed similar interaction profiles, each forming π-anion and π–π stacking interactions with ASP292 and TRP80, hydrogen bonds with SER205, and hydrophobic contacts with LEU210 and LEU264. For the HSP90AA1 protein (PDB: 4U93), luteolin showed the strongest binding energy at −9.90 kcal/mol, forming two hydrogen bonds with ASP93 and TYR139, π–π stacking with PHE138, and hydrophobic interactions with ASN51, LEU107, and THR184. Cryptolepine and apigenin also demonstrated strong affinities of −9.14 and −9.76 kcal/mol respectively, with consistent involvement of PHE138 in π–π stacking and hydrophobic contacts. PHE138 was identified as a conserved hotspot for ligand binding across multiple compounds. Other protein targets such as EGFR, IL-1β, and CASP3 exhibited moderate binding affinities ranging from −7.5 to −8.8 kcal/mol for the selected compounds. Among all tested phytochemicals, cryptolepine consistently displayed the broadest binding spectrum (−6.19 to −10.15 kcal/mol), followed by luteolin and apigenin. In contrast, vernolepin, dihydrovernodalin, and vernomenin generally showed moderate to weak affinities, with fewer instances surpassing the −9.0 kcal/mol threshold (**Table 6**).

**Figure 8.**
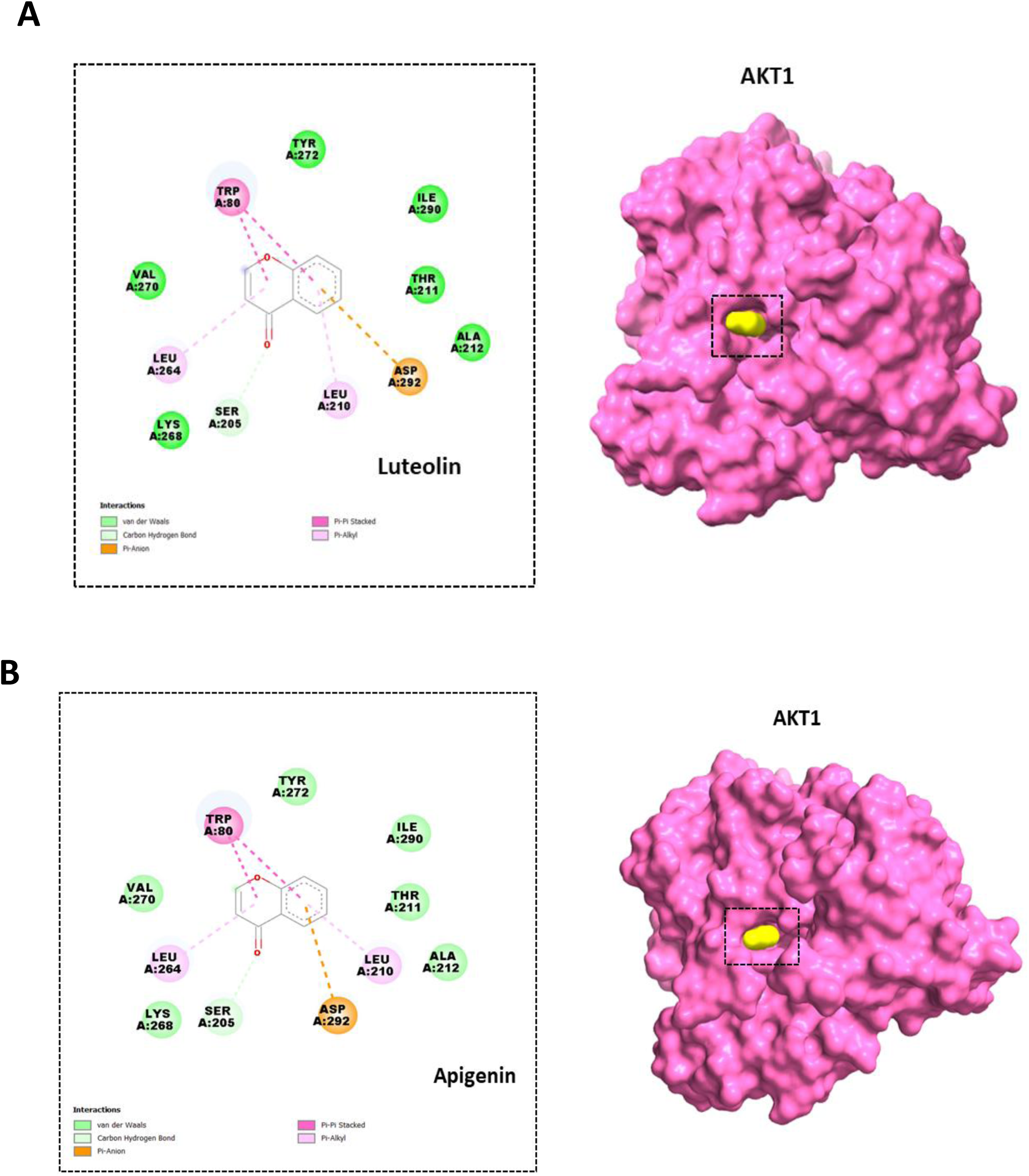
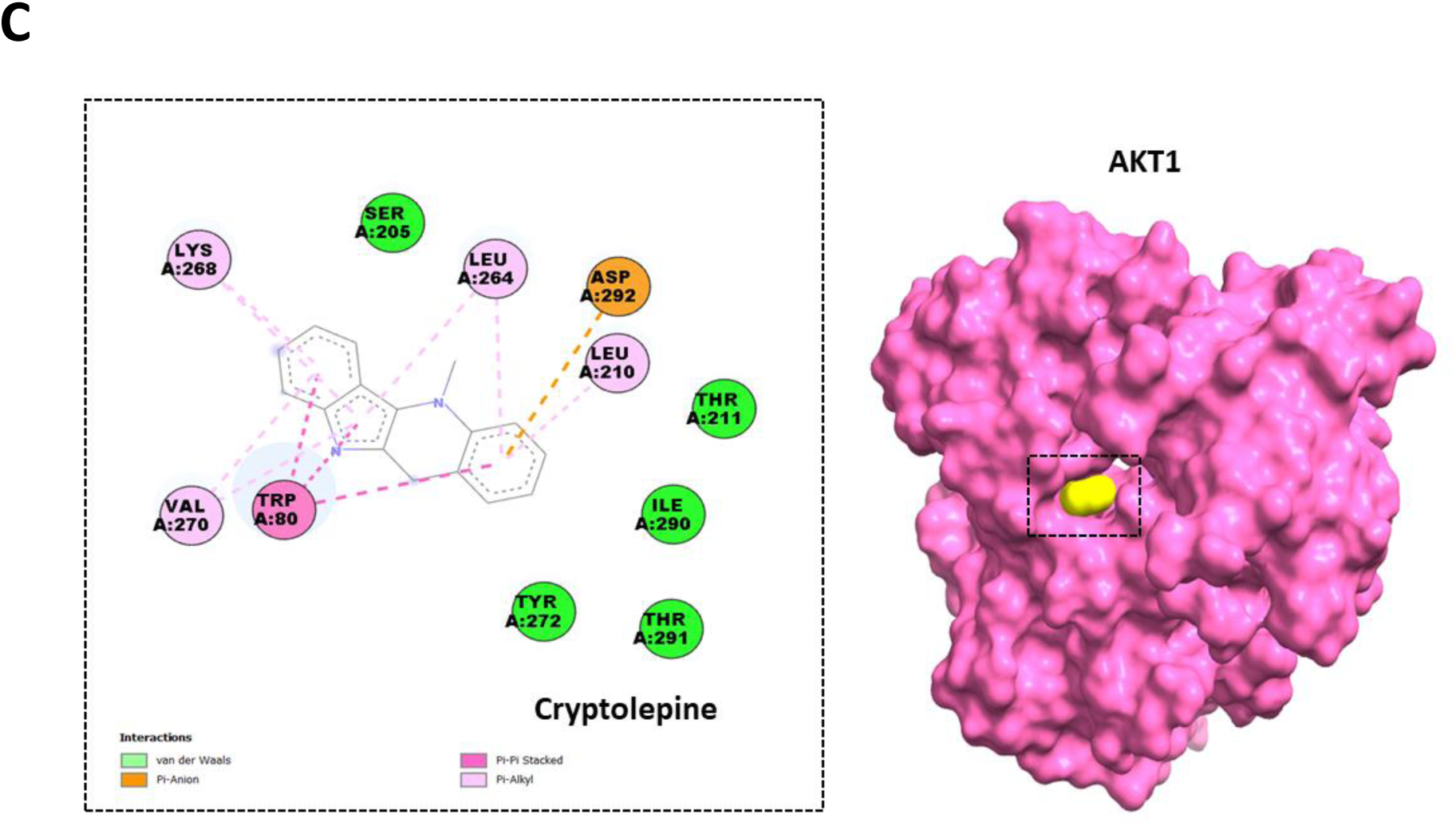
Molecular interactions between the AKT1 protein and the three selected lead compounds: (A) luteolin, (B) apigenin, and (C) cryptolepine.

**Figure 9.**
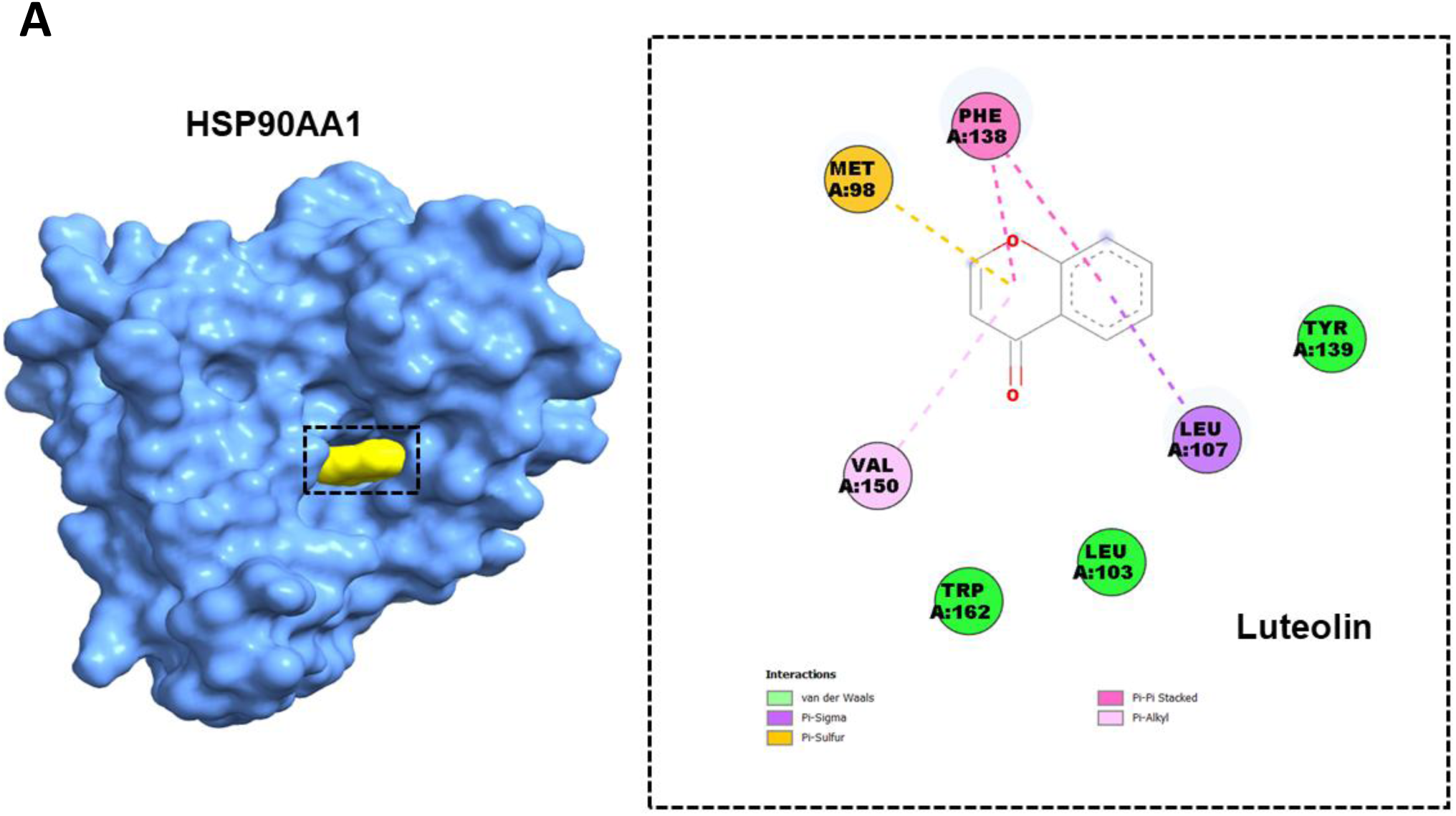
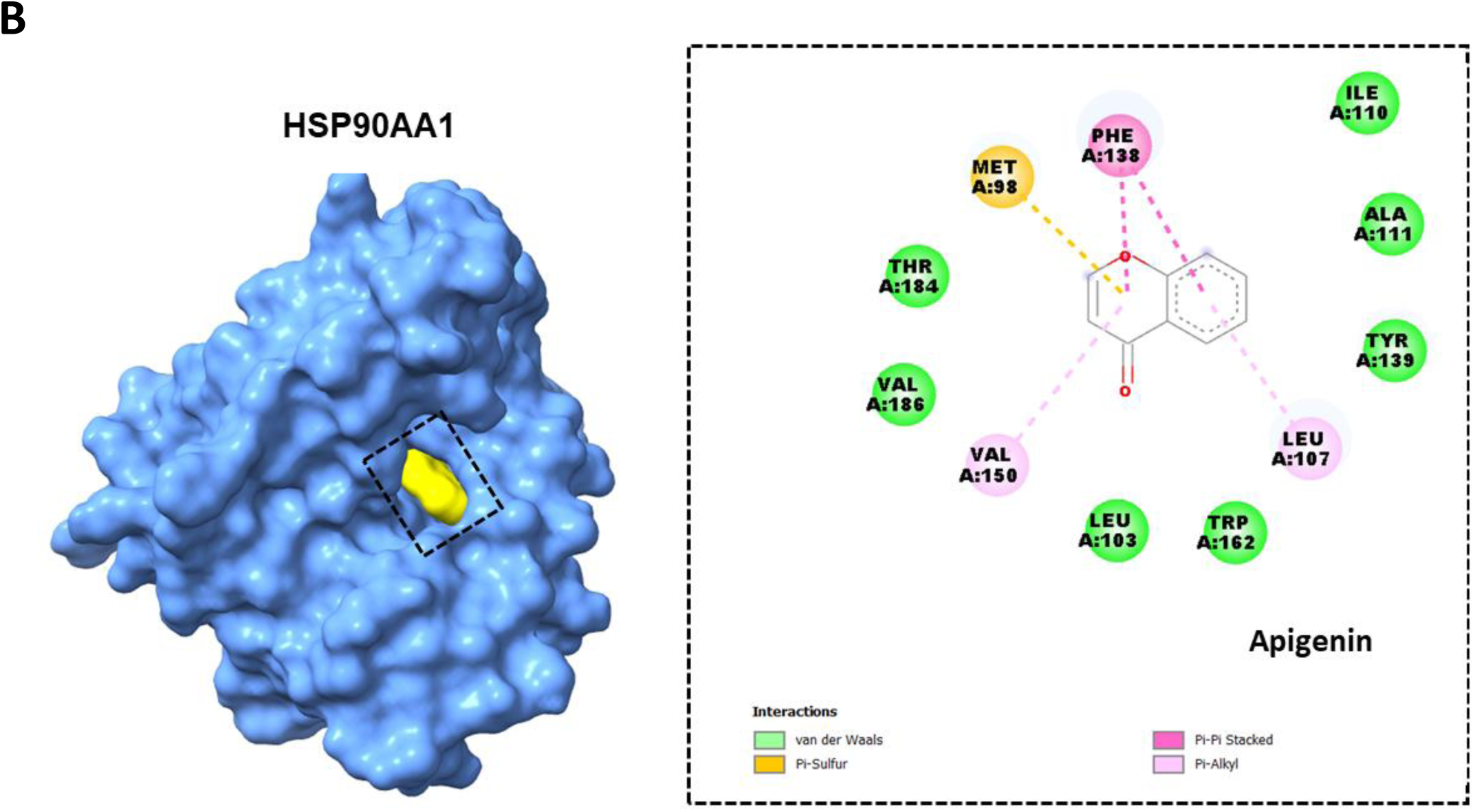
Molecular interactions between the HSP90AA1 protein and the two selected lead compounds: (A) luteolin, and (B) apigenin

**Table 6.**
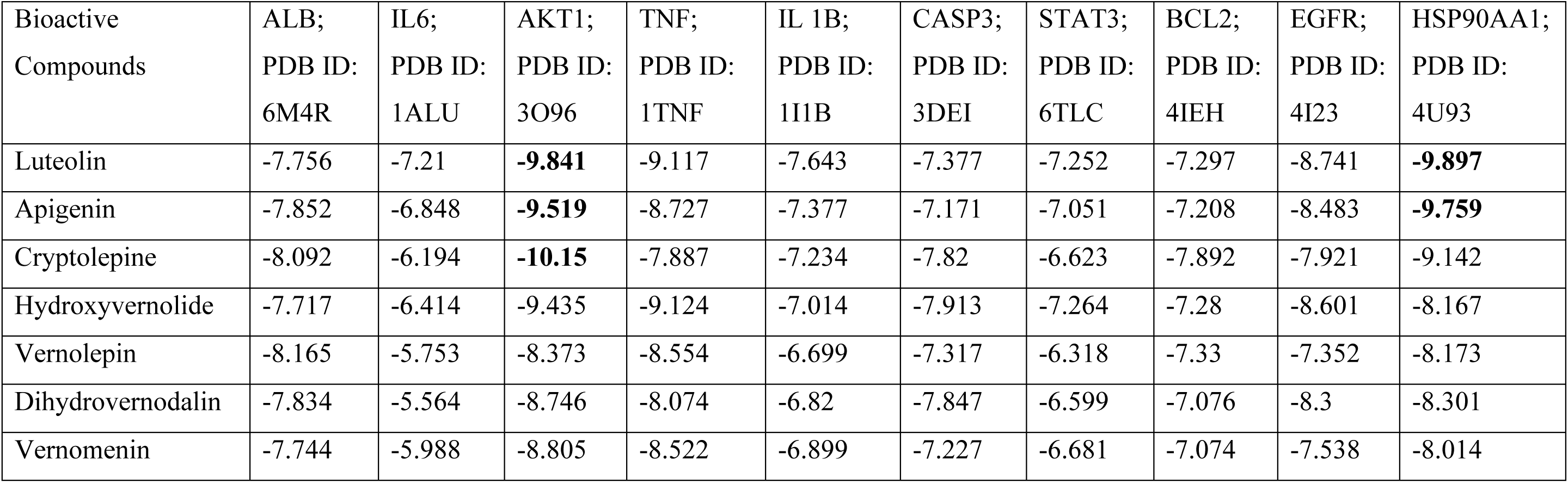
Molecular docking results showing binding energies between 7 bioactive compounds and 10 hub target proteins.

## DISCUSSION

ALD remains a significant global health problem, with chronic ethanol consumption recognized as a major cause of liver inflammation, steatosis, fibrosis, and cirrhosis [**1, 35**]. Phytochemicals are naturally occurring bioactive compounds known for their antioxidant, anti-inflammatory, and hepatoprotective effects [**36**]. This study evaluated the hepatoprotective effects of VA against ethanol-induced liver toxicity in rats using biochemical assays, in silico network pharmacology, and molecular docking studies. Our findings showed that VA significantly improved ALD, possibly through interactions with multiple targets, particularly AKT1 and HSP90AA1. These targets are involved in key cellular processes related to liver injury and repair, including inflammation, oxidative stress, angiogenesis, and metabolic balance.

Phytochemical screening of the VA extract revealed several important secondary metabolites, including saponins, steroids, flavonoids, glycosides, cardiac glycosides, and terpenoids. Phenols, alkaloids, tannins, and carotenoids were absent, suggesting that the hepatoprotective effects of VA may be driven mainly by non-phenolic compounds, although other studies have reported their presence in VA under different conditions **[37]**. These findings align with previous reports highlighting the hepatoprotective and antioxidant potential of VA, especially its flavonoid and saponin content **[38]**. Flavonoids are known to scavenge reactive oxygen species and regulate inflammatory signaling pathways, which are highly involved in ethanol-induced liver injury. Saponins and glycosides contribute to membrane-stabilizing and cholesterol-lowering effects, further supporting their role in liver protection. The combined action of these compounds may explain the biochemical improvements observed in vivo.

Treatment with VA extract significantly reversed the biochemical disturbances caused by ethanol in a dose-dependent manner. It lowered elevated transaminase and bilirubin levels while restoring protein concentrations toward normal ranges, suggesting membrane stabilization and potential hepatocyte regeneration. The highest dose tested (300 mg/kg) produced effects comparable to silymarin, a well-established hepatoprotective agent. The changes observed in liver injury markers such as SGOT (AST), SGPT (ALT), ALKP, total bilirubin, total protein, and albumin reflect both structural damage and functional impairment caused by chronic ethanol exposure. ALT is a more specific indicator of liver injury than AST, which is also present in cardiac and skeletal muscle **[39]**. Elevated ALKP suggests cholestasis or biliary obstruction, possibly resulting from fibrosis or fatty infiltration, while increased total bilirubin points to impaired hepatic clearance and bile excretion **[40]**. Meanwhile, reductions in total protein and albumin indicate compromised synthetic liver function, a hallmark of chronic liver disease **[41]**. Together, these biomarker changes provide a comprehensive picture of the extent and nature of liver damage in this model. These findings indicate that VA potentially exerts its protective effects through multiple mechanisms, including antioxidant activity, anti-inflammatory properties, and preservation of liver cell integrity. The normalization of protein levels may also reflect improved liver synthetic capacity, possibly mediated by enhanced gene expression.

Our results align with previous studies demonstrating the hepatoprotective effects of VA in chemically induced liver injury models. For instance, Iwalokun and colleagues reported that VA reversed acetaminophen-induced elevations in liver enzymes, oxidative stress, and reduced protein levels via antioxidant mechanisms **[10]**. Similarly, Adesanoye and Farombi observed normalization of liver enzymes and increased antioxidant enzyme activity in rats exposed to carbon tetrachloride (CCl₄) **[11]**. Tokofai et al. found decreased liver damage and lipid peroxidation in a broiler chicken model of CCl₄ toxicity following VA treatment **[42]**. Collectively, these studies support the view that VA extract restores liver function possibly by stabilizing cell membranes, reducing oxidative stress, and supporting protein synthesis. However, the precise molecular pathways underlying these effects remain to be clarified, warranting further investigation using transcriptomic or proteomic analyses.

In silico analyses provided insights into the potential multi-target mechanisms of VA in alcoholic liver disease. Seven phytochemicals, including luteolin, apigenin, cryptolepine, vernolide, dihydrovernodalin, hydroxyvernolide, and vernomenin, were found to meet Lipinski’s Rule of Five, indicating good oral bioavailability. This is important in drug development because compounds that do not meet these criteria often have poor absorption and limited therapeutic potential [**34**]. Among these, luteolin, apigenin, and cryptolepine are especially relevant due to their well-known anti-inflammatory, antioxidant, and cytoprotective effects. Sesquiterpene lactones such as vernolide, dihydrovernodalin, hydroxyvernolide, and vernomenin have been less studied but show promising drug-like properties, making them potential novel candidates. Flavonoids like luteolin and apigenin suppress pro-inflammatory cytokines and regulate oxidative stress pathways in liver injury models **[43]**. Cryptolepine, an indoloquinoline alkaloid, is recognized for its antimicrobial and anti-inflammatory activities **[44]**. The presence of these bioactive compounds in VA supports its traditional medicinal use and justifies further investigation as a treatment for liver disease.

Building on this phytochemical profile, molecular docking analyses were conducted to predict how these compounds interact with key protein targets in alcoholic liver disease. The results confirmed the therapeutic potential of VA constituents, particularly luteolin, apigenin, and cryptolepine, which exhibited strong binding to critical ALD-related proteins. Binding energies of –9.5 kcal/mol or lower suggest stable interactions. Cryptolepine and luteolin were identified as top binders for AKT1 and HSP90AA1, respectively. These compounds mainly interact through hydrogen bonds and hydrophobic contacts, which may contribute to their bioactivity. Luteolin’s hepatoprotective effects are well established in vitro and in animal models. In mice treated with DEN and ethanol, luteolin reduced liver inflammation, lipid accumulation, and early cancerous lesions by upregulating SIRT1 and PGC1α expression [**45**]. It also inhibits AKT1, thereby disrupting pro-survival and fibrogenic signaling in liver fibrosis and cancer models [**46**]. In HuH7 cells, luteolin binds AKT1 with high affinity (KD ∼1 µM), modulating oncogenic pathways [**47**]. Furthermore, luteolin is recognized for its potent ability to mitigate liver inflammation in ethanol-induced injury models by suppressing NF-κB activation, restoring AMPK signaling, and preventing lipid accumulation [**48, 49**]. It also attenuates TNF-α-induced activation of NF-κB, MAPK, AKT, JNK, and AP-1 pathways, thereby conferring hepatocyte protection **[50]**.

Docking simulations further revealed that luteolin binds HSP90AA1 through hydrophobic interactions. Inhibiting HSP90AA1 triggers degradation of client proteins such as AKT1 and STAT3, both of which are involved in lipid metabolism, oxidative stress, and inflammation in alcoholic liver disease. Animal models demonstrate that luteolin reduces hepatic steatosis, decreases pro-inflammatory cytokines, and restores antioxidant enzyme activity [**51**]. Given that ALD involves oxidative stress, inflammation, and abnormal activation of survival pathways such as PI3K/AKT and MAPK, targeting HSP90AA1 represents a promising multi-target strategy. Luteolin binding may suppress inflammation, inhibit fibrogenic signaling, and promote apoptosis of damaged hepatocytes and activated stellate cells. This aligns with previous findings that HSP90 inhibition reduces liver injury in preclinical models [**52**]. Overall, these results suggest that luteolin functions both as an antioxidant and as a regulator of protein stability, supporting emerging interest in HSP90 inhibitors as therapeutic agents.

We found that apigenin also demonstrated strong binding to AKT1 and HSP90AA1, sharing similar binding residues as luteolin, however, with distinct bond lengths. Structurally, apigenin is similar to luteolin but lacks a hydroxyl group at the 3′ position of the B-ring. Just as luteolin, apigenin exhibits hepatoprotective effects by modulating inflammation, reducing oxidative stress, and inhibiting fibrogenesis in both in vitro and in vivo models. It reduces CYP2E1 activity, a major source of reactive oxygen species during ethanol metabolism, thereby limiting oxidative damage and hepatic fat accumulation [**53**]. Additionally, it enhances antioxidant defenses and improves lipid metabolism through activation of PPARα **[53]**. Apigenin’s anti-inflammatory activity is partly mediated through inhibition of the NLRP3/NF-κB pathway, which drives chronic inflammation by promoting cytokine production, including TNF-α, IL-6, and IL-1β [**54**]. It also suppresses hepatic stellate cell activation, leading to lower ALT and AST levels and reduced extracellular matrix deposition **[55]**. Importantly, apigenin inhibits the PI3K/AKT/mTOR pathway, reducing cell proliferation and promoting apoptosis, as shown in liver injury and cancer models [**56**]. Our docking analysis revealed that apigenin binds to an allosteric site on AKT1 involving key residues critical for allosteric inhibition. Structural studies support the functional relevance of this pocket **[57]**, suggesting apigenin may act as a natural allosteric modulator. This mechanism likely underlies its therapeutic effects in ALD and warrants further investigation. Apart from, AKT1, apigenin also binds to the N-terminal ATP-binding site of HSP90AA1, a well-characterized ligand-binding region [**58**]. This interaction may disrupt the HSP90–Cdc37 complex, promoting degradation of client proteins such as AKT and STAT3, thereby attenuating inflammatory and oncogenic signaling **[59]**. In the context of ALD, apigenin acts on multiple interconnected pathways, including antioxidant regulation (Nrf2), inflammation (NF-κB, NLRP3), metabolism (AMPK/SREBP, PPARα/γ), fibrosis (TGF-β1/Smad3, Wnt/β-catenin), and cell survival/apoptosis (PI3K/AKT/mTOR, MAPKs, caspases) [**48**]. Our findings are consistent with earlier studies identifying AKT1 and HSP90AA1 as key targets of apigenin, luteolin, and cryptolepine in ALD [**60**]. Dual inhibition of these proteins may provide synergistic therapeutic effects by simultaneously disrupting survival signaling and stress responses. Although apigenin exhibits low toxicity, its clinical application is limited by poor oral bioavailability, underscoring the need for optimized delivery systems and further safety evaluation [**61**].

Cryptolepine, an indoloquinoline alkaloid from *Cryptolepis sanguinolenta*, also interacted with the allosteric pocket of AKT1, contacting residues targeted by known inhibitors. While direct evidence for cryptolepine’s effects in ALD is limited, related alkaloids have been shown to modulate kinase signaling, induce apoptosis, and disrupt protein stabilization by chaperones in other disease models [**62**]. These findings suggest a potential therapeutic role for cryptolepine, although further experimental work is needed to confirm its effects on AKT1 phosphorylation and HSP90 client protein stability. Future studies should evaluate whether this dual-target mechanism can improve outcomes in preclinical ALD models.

GO and KEGG pathway enrichment analyses provided functional implications to the 173 subnetwork targets identified from the top 10 hub genes. GO analysis revealed significant enrichment in biological processes, including responses to nitrogen compounds, oxidative stress, regulation of protein phosphorylation, and small molecule metabolism. These processes are central to ALD pathophysiology. Chronic alcohol intake increases reactive nitrogen and oxygen species, causing oxidative and nitrosative stress that damages hepatocytes and promotes lipid peroxidation **[63]**. The enrichment of oxidative stress processes aligns with the antioxidant effects of flavonoids such as luteolin and apigenin, which increase the activity of enzymes, including superoxide dismutase and catalase, thereby mitigating liver injury [**64**]. Phosphorylation processes indicate that VA compounds may modulate kinase signaling pathways involved in cell survival, inflammation, and tissue repair [**65]**. This aligns with our docking results, which demonstrate strong predicted binding to key kinases such as AKT1 and regulatory proteins like STAT3.

In the molecular function category, targets were enriched for ATP binding, protein kinase activity, and nucleotide binding. These functions are essential for energy-dependent enzymatic activity and cellular signaling, both of which are often impaired in ALD due to mitochondrial dysfunction and ethanol-induced metabolic imbalance [**66**]. This supports the hypothesis that VA compounds interact with enzymes and signaling proteins involved in energy metabolism and stress responses, potentially contributing to the restoration of hepatocellular homeostasis. In this study, cellular component enrichment highlighted membrane rafts, extracellular vesicles, and synaptic membranes as key sites of action. These structures are important for signal transduction and intercellular communication, particularly in inflammatory liver disease [**67**]. Ethanol alters vesicle signaling and exosome release, facilitating immune cell recruitment and fibrogenesis [**68**]. Therefore, the predicted effects of VA compounds on these components may contribute to their multi-target potential, especially in modulating liver cell communication and immune responses.

Furthermore, KEGG pathway analysis identified 222 significantly enriched pathways, including PI3K–Akt, VEGF, HIF-1, TNF, IL-17, and FoxO signaling. These pathways regulate inflammation, angiogenesis, oxidative stress responses, and cell survival, all of which are disrupted during ALD progression [**69**]. For example, the PI3K–Akt pathway promotes hepatocyte survival and liver regeneration [**70**]. Flavonoids such as luteolin and apigenin have been reported to activate or stabilize this pathway, consistent with their high affinity for AKT1 observed in this study. Moreover, HIF-1 signaling is upregulated in hypoxic liver tissue, driving angiogenesis and metabolic adaptation. VA compounds may aid tissue repair by targeting this pathway [**71**]. Additionally, pathways related to lipid metabolism and atherosclerosis were also enriched, suggesting broader metabolic benefits. Alcohol-induced steatosis results from impaired lipid storage and fatty acid oxidation. VA compounds may counteract this by activating AMP-activated protein kinase and downregulating sterol regulatory element-binding proteins, thereby reducing hepatic lipid accumulation [**72**].

## CONCLUSION

In this study, the hepatoprotective potential of VA leaf extract against ethanol-induced liver injury was confirmed. The in vivo findings demonstrated significant restoration of liver function biomarkers, while the in-silico analyses revealed that flavonoids and sesquiterpene lactones in the extract interact with key proteins involved in oxidative stress, inflammation, and apoptosis, particularly AKT1 and HSP90. Furthermore, these compounds exhibited favorable pharmacokinetic and safety profiles, reinforcing their drug-likeness and therapeutic promise. Overall, VA emerges as a promising candidate for the development of plant-based interventions for alcoholic liver disease. Future research should focus on isolating the active constituents and validating their efficacy in both preclinical and clinical settings.

